# Hotspot coevolution at protein-protein interfaces is a key identifier of native protein complexes

**DOI:** 10.1101/698233

**Authors:** Sambit K. Mishra, Sarah J. Cooper, Jerry M. Parks, Julie C. Mitchell

## Abstract

Protein-protein interactions play a key role in mediating numerous biological functions, with more than half the proteins in living organisms existing as either homo- or hetero-oligomeric assemblies. Protein subunits that form oligomers minimize the free energy of the complex, but exhaustive computational search-based docking methods have not comprehensively addressed the protein docking challenge of distinguishing a natively bound complex from non-native forms. In this study, we propose a scoring function, *KFC-E*, that accounts for both conservation and coevolution of putative binding hotspot residues at protein-protein interfaces. For a benchmark set of 53 bound complexes, *KFC-E* identifies a near-native binding mode as the top-scoring pose in 38% and in the top 5 in 55% of the complexes. For a set of 17 unbound complexes, KFC-E identifies a near-native pose in the top 10 ranked poses in more than 50% of the cases. By contrast, a scoring function that incorporates information on coevolution at predicted non-hotspots performs poorly by comparison. Our study highlights the importance of coevolution at hotspot residues in forming natively bound complexes and suggests a novel approach for coevolutionary scoring in protein docking.

**Author Summary:** A fundamental problem in biology is to distinguish between the native and non-native bound forms of protein-protein complexes. Experimental methods are often used to detect the native bound forms of proteins but, are demanding in terms of time and resources. Computational approaches have proven to be a useful alternative; they sample the different binding configurations for a pair of interacting proteins and then use an heuristic or physical model to score them. In this study we propose a new scoring approach, *KFC-E*, which focuses on the evolutionary contributions from a subset of key interface residues (hotspots) to identify native bound complexes. *KFC-E* capitalizes on the wealth of information in protein sequence databases by incorporating residue-level conservation and coevolution of putative binding hotspots. As hotspot residues mediate the binding energetics of protein-protein interactions, we hypothesize that the knowledge of putative hotspots coupled with their evolutionary information should be helpful in the identification of native bound protein-protein complexes.

## Introduction

Binding interactions between proteins regulate life at a molecular level. Such interactions can lead to the creation of new functional interfaces in bound complexes, which can act as active sites or can lead to increased thermal stability of the complex [1,2] compared to the isolated subunits. Moreover, the dynamics of the individual binding partners may be altered [3], allowing for new conformational ensembles to be sampled. Consequently, an important outcome of such interactions is the ability of a complex to orchestrate novel molecular functions often not possible by the monomeric forms [4]. However, proteins often bind in specific conformations, commonly referred to as native state conformations, which are energetically favorable and allow the complex to execute its function.

Experimental methods that probe the native bound state of interacting proteins range from assays that inform only whether proteins interact or not, such as yeast two-hybrid assays and co-immunoprecipitation, to structural determination methods such as nuclear magnetic resonance (NMR) spectroscopy, X-ray crystallography and cryo-electron microscopy (cryo-EM). Numerous computational approaches [5] have also been developed to address the CAPRI (Critical Assessment of PRedicted Interactions) [6] protein-protein docking challenge, an effort aimed at identifying the native bound form of interacting proteins. Simply put, the goal of this challenge is to identify the energetically most favorable docked conformations between two or more proteins. Existing rigid and flexible docking protocols [7] initially perform exhaustive sampling to obtain a wide range of binding modes for interacting proteins. These configurations are then assessed using a scoring function that ranks the predicted binding poses from most to least favorable. The binding conformation, or cluster of conformations, with the most favorable binding score is predicted to be the native binding mode.

The scoring function is the key component of any docking method. Historically, scoring functions have accounted for the electrostatic interactions, desolvation energies, van der Waals interactions and the extent of complementarity of the interfaces of the binding partners [8,9]. However, more recent methods also incorporate the knowledge of interfacial residues [10] and evolutionary information of these residues [11] into their scoring functions to guide protein-protein docking. These approaches are inspired by work suggesting that interfacial residues tend to be under higher evolutionary selection pressure than non-interfacial residues, resulting in either high conservation [12,13] or strong coevolution [14] with the interacting residues in the binding partner. Methods such as EVcomplex [15], GREMLIN [16], and DCA [17] use the coevolution of interfacial residues of the binding partners as a guide to identify binding modes, whereas InterEvScore [18] uses both conservation and coevolution. Coevolution analysis has also been used to detect allosteric pathways [19], suggesting a strong evolutionary linkage between distant allosterically coupled residues, as well as, for *de novo* structure prediction of proteins [20].

The binding interactions between subunits of non-transient protein complexes are often strongly mediated by a subset of interface residues, often referred to as hotspots [21]. Hotspot residues contribute significantly to the binding energetics of a protein complex and can be identified experimentally through alanine scanning mutagenesis, i.e., substituting each residue at the interface with alanine and measuring the corresponding binding free energy change relative to the wild type (ΔΔ*G_bind_*) for the entire complex. Mutation of hotspot residues to alanine results in a larger change in the binding free energy of the complex (≥2 kcal/mol) than mutation of non-hotspot residues [21,22]. Such mutagenesis experiments, however, are laborious and time consuming. Thus, a number of computational initiatives have been undertaken toward computational hotspot prediction [22–28]. Owing to their key role in mediating protein-protein interactions, prior knowledge of hotspot residues has also been employed in protein-protein docking [29]. In a reported study, a library of potential hotspot residues and their rotamers were used to produce high-affinity binding conformers for influenza hemagglutinin [30]. Owing to their propensities to be involved in strong interactions, hotspot residues were suggested to have fewer conformational degrees of freedom and their knowledge restricted the number of binding conformations, facilitating the identification of high-resolution binding modes. Another study used hotspot residues predicted with the KFC2 (Knowledge-based FADE and Contacts) method [28] to identify near-native binding modes. KFC2 uses features that are based on residue solvent accessibilities, flexibilities and atomic densities within a machine-learned framework to predict hotspot residues with high confidence. Predicted hotspots from KFC2 were previously coupled with calculations of absolute and chemical conservations of the hotspots and their contacts in different scoring schemes to identify natively bound poses [31]. For a small dataset of 17 complexes, using one of the scoring schemes led to precise identification of near-native poses for more than 80% of the complexes. These studies demonstrate that predicted hotspot residues can be beneficial in identifying natively docked conformations of protein complexes.

Within protein interfaces, we expect that hotspot residues are more highly conserved than non-hotspot residues. However, in cases in where interacting partners have evolved, coevolution is observed at the interface to maintain the interactions between the binding partners [14]. In such cases, coevolution analysis is expected to be a better predictor of near-native models of protein-protein complexes than conservation. Thus, a combined scoring function incorporating both conservation and coevolution should in principle, be more accurate in identifying near-native poses.

Here we report *KFC-E*, a scoring function that is based on both the coevolution and conservation of binding hotspots at protein-protein interfaces. While previous work has utilized coevolutionary data to set sampling restraints for docking, we are unaware of other work that utilize coevolution at hot spots for scoring docked configurations. The binding hotspots for our method were predicted with KFC2 [28]. To verify the effect of including both coevolution and conservation, we compare the performance of *KFC-E* in terms of identifying near-native poses with scoring functions that use either residue conservation or coevolution or neither. We also verify the importance of including the information on hotspot coevolution by verifying the performance with respect to a scoring function using non-hotspot coevolution. We test these scoring functions on a benchmark set of 53 bound and 17 unbound protein complexes for which the binding modes were exhaustively sampled with ZDOCK [9]. Our findings stress the importance of conservation and coevolution of hotspot residues in maintaining native interfaces over the course of evolution.

## Materials and Methods

### Dataset of Bound and Unbound Structures

We utilized a subset of 53 bound and 17 unbound heterodimeric proteins from a benchmark set of protein-protein complexes [16]. For the unbound set, we used an available apo structure or apo structure of a close homolog in place of one of the bound chains. HHpred [32] was then used to align the bound sequences to the sequences of the unbound templates for threading, followed by homology modeling with RosettaCM [33] to generate 1,000 models of each protein. For each case, we selected the top-scoring model and superimposed it onto the bound-bound complex for protein-protein docking. Tables 1 and 2 report the PDB IDs of the complexes included for the bound and unbound datasets, respectively.

### Protein-Protein Docking and Interface RMSD Calculations

We used ZDOCK [9,34] to perform rigid docking with exhaustive and dense sampling, resulting in 54,000 docked poses for each complex. For each complex, we then identified interface residues as those with a heavy atom within 5 Å of a heavy atom of any residue in the binding partner. We calculated the interface root-mean squared deviation (i-RMSD) [35] for a given docked pose as the RMSD between the Cα atoms of the interfacial residues in the native complex and the docked pose.

### Hotspot Prediction

We used KFC2 [28] to predict hotspots for each binding pose sampled by ZDOCK. KFC2 comprises two prediction methods: KFC2a and KFC2b, which each use different subsets of features to predict hotspot residues. For our calculations, we use the predictions from KFC2a as it has higher predictive accuracy than KFC2b.

### Residue-Residue Coevolution

The benchmark dataset [16] also includes paired multiple sequence alignments for each protein complex. We used these alignments as input to CCMpred [36] with default parameters to generate a coevolution matrix and estimate the extent of coevolution between all residue pairs in the complex.

### Residue Conservation

To calculate the extent of conservation at each residue position in a given complex, we randomly sampled 150 sequences from the paired alignment of each complex. For each complex, we performed 50 repetitions of random sampling to obtain 50 subsets of aligned orthologs. Using rate4site [37] with default parameters, we calculated the conservation scores for each sampled sequence set and then the average conservation score for each residue over the 50 subsets of sequences. Rate4site reports the extent of evolution for each residue as normalized scores in which a negative score implies a slower rate of evolution and thus, higher conservation. In our calculations, we first inverted the scale on the reported scores so that the more conserved residues have more positive scores and then shifted the inverted scores by adding the absolute value of the minimum of the inverted scores to each score. Following these operations, the least conserved residue had a score of 0 and the more conserved ones had positive scores.

### Scoring Functions

We used eight different scoring functions (Table 3) to rank the binding poses sampled by ZDOCK. Each scoring function is described in detail below.

#### a. HC Co-evol: Coevolution of hotspot residues with contacts in the binding partner

For each subunit in a given binding pose, we consider the extent of coevolution between the hotspot residues predicted by KFC2a and their contacts in the binding partner. In this context, we define a residue *r*_*j*_ in chain B to be in contact with a predicted hotspot residue *r*_*i*_ in chain A if any of the heavy atoms of residue *r*_*j*_ is within 7 Å of any heavy atom of *r*_*i*_. If *hotspots*_*A*_ and *hotspo ts*_*B*_ are sets of hotspots predicted by KFC2a in chains A and B, respectively, and *contacts*_*B*_ and *contacts*_*A*_ are their contacts in the respective binding partner, then the two chains from a given binding pose *n* are scored as follows.

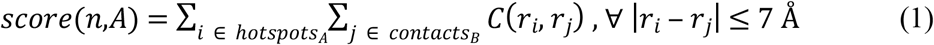

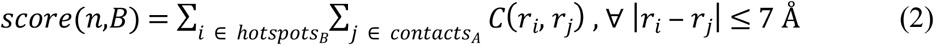

where *score*(*n*,*A*) is the score for pose *n* from chain *A* and *score*(*n*,*B*) is the score for pose *n* from chain *B*. *C* is the coevolution matrix for the complex obtained with CCMpred. The summation is over all pairs of hotspot residues and their contacts whose Euclidean norm |*r*_*i*_ ‒ *r*_*j*_| is within 7 Å. The scores obtained for chains A and B of each pose from the sampled set of 54,000 binding poses are stored in two separate vectors: *S*_*A*_ and *S*_*B*_. Thus, the *i*th element in *S*_*A*_ and *S*_*B*_ corresponds to *score*(*n*,*A*) and *score*(*n*,*B*) of a given pose *n*, respectively, and the dimensions of both *S*_*A*_ and *S*_*B*_ equals the total number of sampled binding poses, i.e., 54,000.

To obtain a cumulative score for a given pose, we then perform element-wise addition of the vectors *S*_*A*_ and *S*_*B*_ as follows:

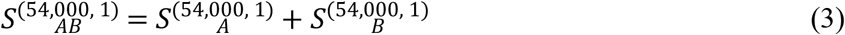

and normalize *S*_*AB*_ to range between 0 and 1 with the following equation:

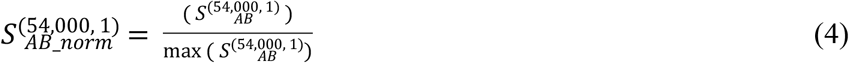

Here, 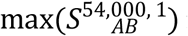 is the maximum score in *S*_*AB*_. The scores of all binding poses thus range from 0 to 1, with the most probable near-native pose having a score of 1. Poses for which there are no predicted hotspots by KFC2a are assigned a score of 0. This step of obtaining a single normalized score for each pose is the same for all the scoring functions used in this study.

#### b. HH Co-evol: Coevolution of hotspot residues with hotspot residues in the binding partner

Instead of considering the coevolution between hotspots and all their contacts across the protein interface (as done for *HC Co-evol*), here we consider for a given subunit in a binding pose the extent of coevolution with contacts that are also hotspots. The scoring function for a given pose *n* is described in Eq. 5.

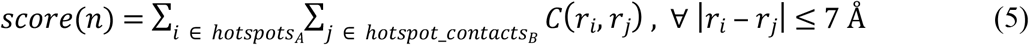

where *hotspots*_*A*_ is the set of predicted hotspots on chain A and *hotspot*_*contacts*_*B*_ is the set of hotspots in chain B that are in contact with the hotspots in chain A. Unlike the previous scoring function, we do not score both chains because the hotspot-hotspot contact pairs for chain A are identical to that for chain B.

#### c. NH-NH Co-evol: Coevolution of non-hotspot residues with non-hotspot residues in the binding partner

Here we consider all residues that are not predicted as hotspots by KFC2a (i.e., non-hotspots) and their extent of coevolution with non-hotspot residues in the binding partner. Like the *HH Co-evol* scoring function, the contact pairs for each chain are the same and thus, we score a given pose *n* based on the contacts from a single chain as indicated in Eq. 6.

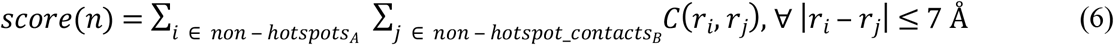

Here, *non* ‒ *hotspots*_*A*_ and *non* ‒ *hotspot*_*contacts*_*B*_ are the non-hotspots in chain A and the corresponding non-hotspot contacts in chain B, respectively.

#### d. HC Cons: Conservation of hotspot residues and their contacts in the binding partner

This scoring function is similar to *HC Co-evol* but instead of using the extent of coevolution between hotspots and their contacts in the binding partner we use the residue-level conservation scores (*E* (*r*_*i*_, *r*_*j*_)) from rate4site to score the hotspots and their contacts.

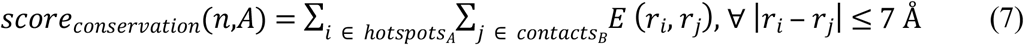

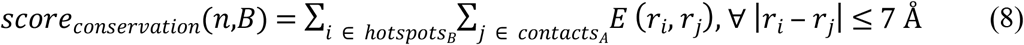

Then, we use an approach similar to the *HC Co-evol* scoring function to obtain a cumulative score for a given pose using the scores from each chain.

#### e. Num H: Number of hotspots

A given pose is scored by the total number of predicted hotspots on the two chains.

#### f. Num HC: Number of hotspot contacts

We consider the predicted hotspots on a given chain for a given pose and the number of residues in the binding partner in contact with the predicted hotspots. This approach is similar to the *HC Co-evol* scoring function, except that we do not consider the extent of coevolution between the hotspots and their contacts in the binding partner.

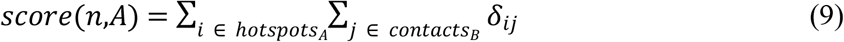

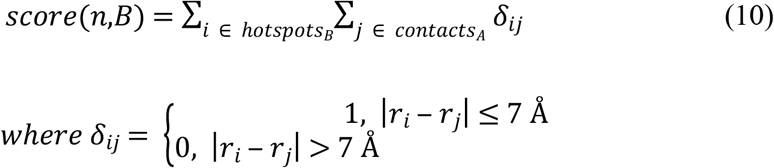

A given pose is then scored using the approach described for *HC Co-evol* to obtain a cumulative score between 0 and 1.

#### g. Num IC: Number of interface contacts

We score each pose based on the total number of unique residue-residue contacts across the interface, which are identified using a distance cutoff of 7 Å.

#### h. KFC-E: Joint coevolution and conservation function

To score a binding mode, we consider not only the coevolution between hotspots and their contacts in the binding partner, but also the extent of conservation of the hotspot residues and the extent of coevolution between the contacts for a given hotspot residue (see Fig. 2). Equations 11 and 12 describe the scoring scheme.

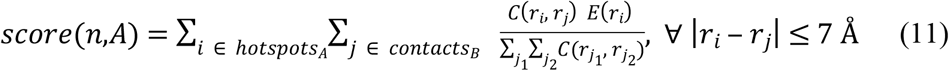

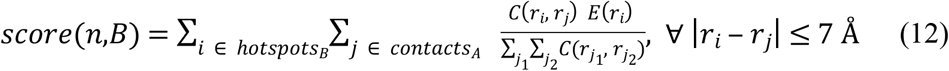

**Figure 1.**
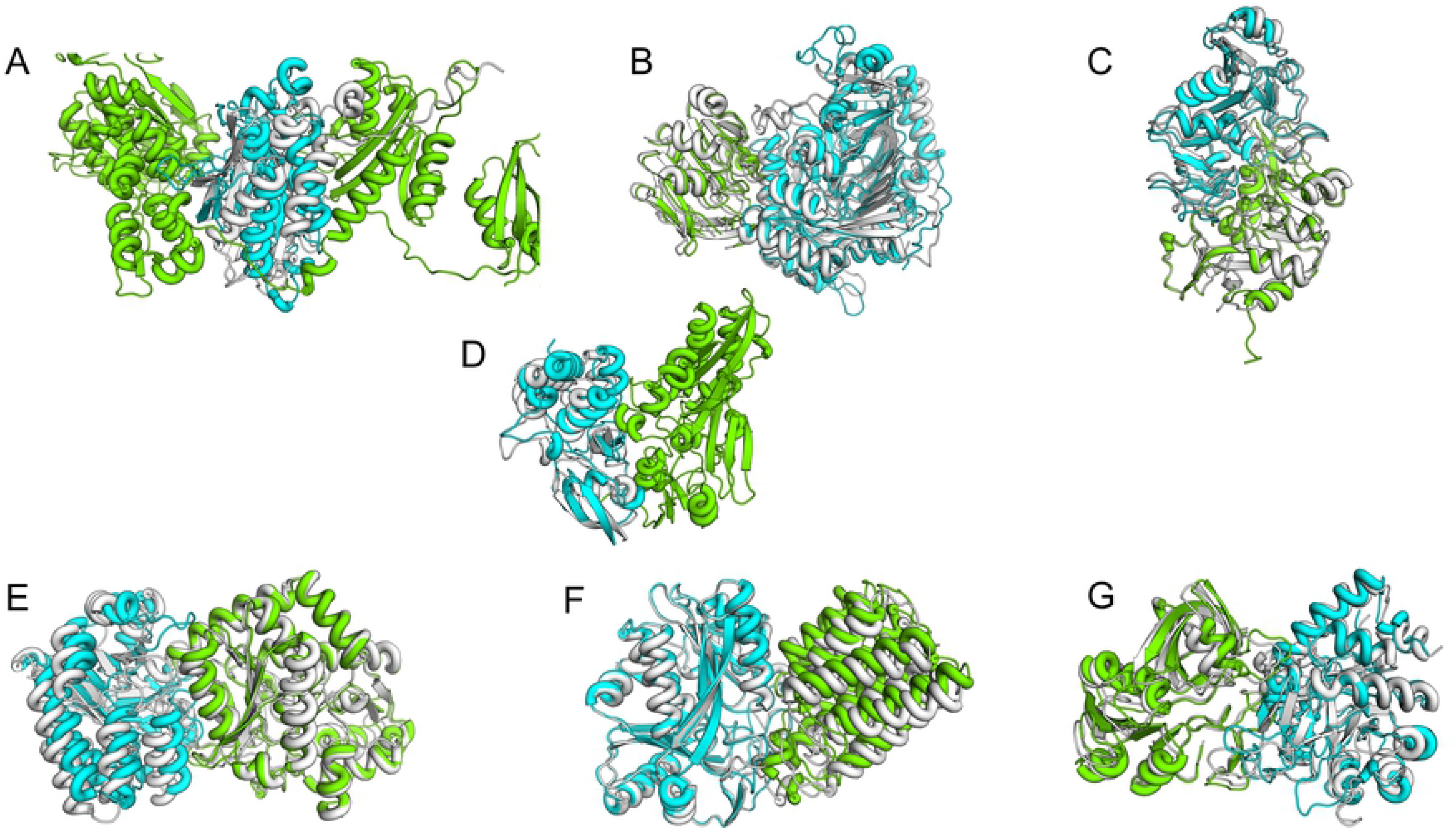
Trends for the *HC Co-evol*, *HH Co-evol*, *NH-NH Co-evol* and *HC Cons* scoring functions for three complexes: dihydroorotate dehydrogenase (PDB entry 1EP3, PyrD and PykR subunits), tryptophan synthase (PDB entry 1QOP, alpha and beta chains) and toxin-antitoxin complex RelBE2 (PDB entry 3G5O, chains A and B). Each column shows the scoring trends for a specific scoring function on all the three complexes. Each row shows the trends across the four scoring functions for a given complex.

**Figure 2.**
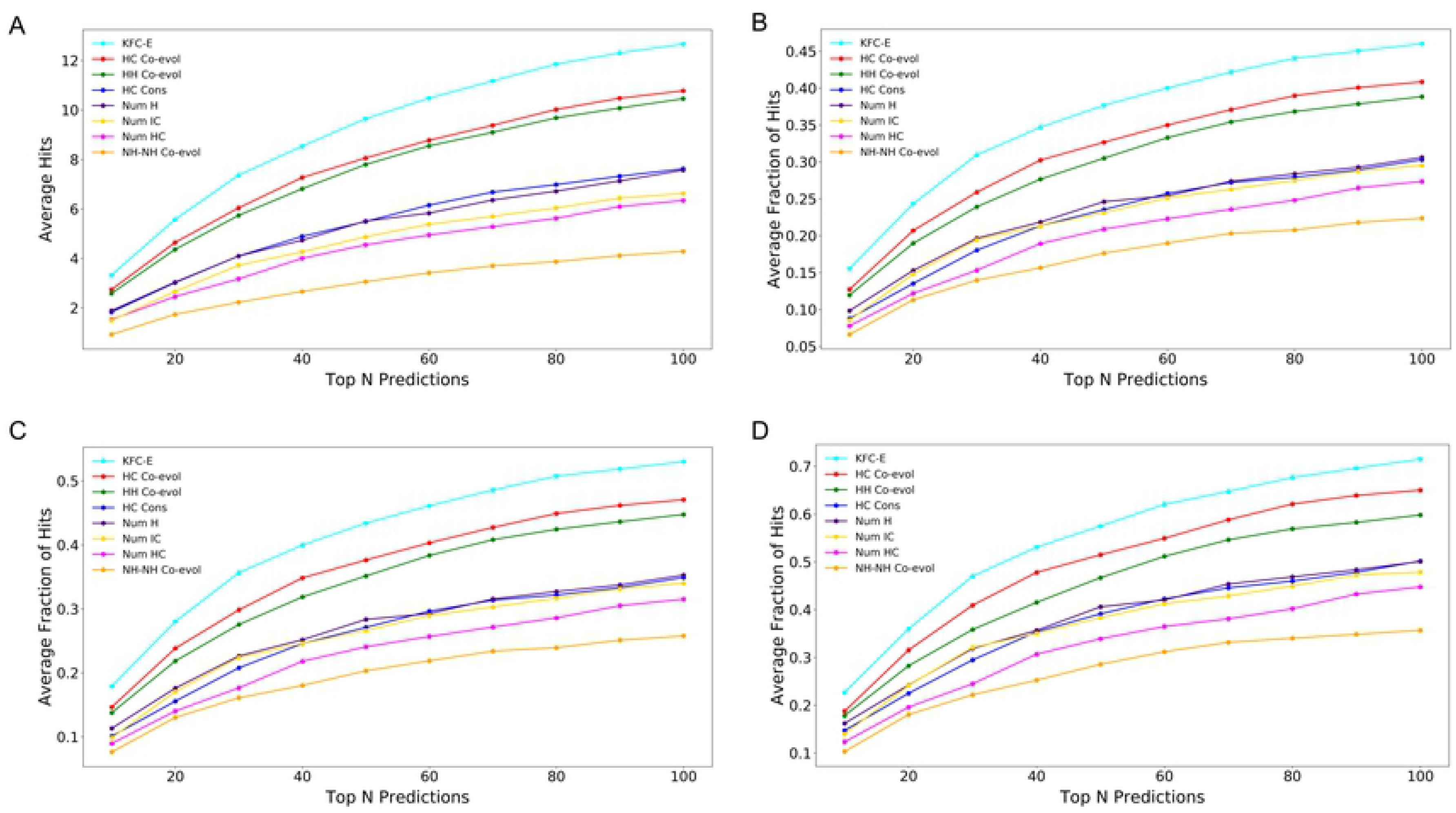
Inter- and intra-protein coevolution in *KFC-E*. Hotspot residues on chain A (red spheres) and their contacts on chain B (gray spheres) are shown. Two types of coevolutionary information taken into account by the *KFC-E* scoring function are shown: (**A**) Coevolution between hotspots and their contacts across the interface (blue arrows) and (**B**) coevolution among the contacts at the interface of chain B (yellow arrows).

The term in the denominator ∑_*j*_ ∑_*j*_ *C*(*r*_*j*1_, *r*_*j*2_) accounts for the cumulative coevolution between all contacts across the interface for a given hotspot residue *r*_*i*_, while *E*(*r*_*i*_) is the extent of conservation reported by rate4site for hotspot residue *r*_*i*_. If the number of contacts for a given hotspot residue is less than 2, we set the denominator to 1.

## Results and Discussion

We utilized a set of 53 bound and 17 unbound bacterial protein-protein complexes (Tables 1 and 2) taken from a previously benchmarked dataset [16], with an aim to test the performance of our new scoring function for protein-protein docking, *KFC-E*. In this study we address the problem of scoring docked poses that have already been generated by exhaustive search docking software rather than sampling of the poses. Our underlying assumption is that incorporating the knowledge of highly probable hotspot residues into a scoring framework that uses both evolutionary and coevolutionary information of these residues should, in principle, facilitate the identification of natively docked poses. Because hotspot residues strongly influence the binding energetics between protein-protein interactions, they experience strong evolutionary selection. Therefore, these residues tend to be evolutionarily more conserved than other residues at the interface of protein-protein complexes. However, in cases where both binding partners evolve significantly [14], hotspot residues may be less conserved, but should coevolve strongly with residue contacts in their binding partner in order to maintain inter-protein interactions in a given complex. Thus, incorporating both conservation and coevolution of hotspots and their contacts in a joint scoring function should in principle provide higher precision in identifying native-like binding modes than using either conservation or coevolution alone.

The present study requires the both evolutionary and coevolutionary information. For this purpose, we used previously generated paired multiple sequence alignments of potentially interacting orthologs [16] for the protein complexes in the dataset. For the calculation of residue-residue coevolution it is essential to filter out paralogs while maintaining high confidence in the interactions among the paired orthologs. Thus, if A and B are the interacting subunits in a complex and A′ is an ortholog of A, while B′ and B′′ are orthologs and paralogs of B, respectively, then A′ must be paired with B′ and not with B′′. This is a non-trivial problem for eukaryotes that have multiple paralogs. In prokaryotes, however, genes encoding interacting proteins are often co-localized in operons and thus, it is simpler to generate paired alignments of orthologs for prokaryotes than for eukaryotes. Using paired alignments, we calculated residue-residue contact maps for each complex and then performed multiple iterations of random sampling (described in Materials and Methods) to calculate the average residue-level conservation. We performed exhaustive and dense sampling of binding modes for each complex in the bound and unbound datasets and then predicted hotspots for each pose. We then verified the performance in scoring the docked poses using eight different scoring functions, each incorporating information about a different aspect of the binding interface to score the poses. Our goal was to identify the scoring scheme that shows the best performance in terms of discriminating near-native complexes from non-native ones. A summary of the different scoring functions is provided in Table 3 and a more detailed description is given in the Materials and Methods.

We first assessed the performance of four scoring functions that use either coevolution or conservation of the binding interface for the complexes in the bound dataset (Table 3). *HC Co-evol* considers the coevolution of hotspots with all contacting residues in the binding partner, *HH Co-evol* considers the extent of coevolution of hotspot residues with contacting hotspot residues in the binding partner, *NH-NH Co-evol* considers coevolution of non-hotspot residues with non-hotspot contacts in the binding partner, and *HC Cons* considers the extent of conservation of hotspots and their contacts in the binding partner. We then introduced the joint scoring function (*KFC-E*), which incorporates both coevolution and conservation, and assessed its performance in comparison to the above four scoring functions. Third, we considered three additional scoring functions that do not take into consideration any evolutionary or coevolutionary information of the interface residues. Instead, for a given binding pose these scoring functions compute either the total number of predicted hotspots (*Num H*), or the total number of contacts for the predicted hotspots (*Num HC*) or just the number of contacts in the binding interface (*Num IC*). We then determined which of the eight scoring functions performed best for the proteins in the bound dataset. Lastly, we verify the performance of this scoring function for the complexes in the unbound dataset and address some of the more general characteristics and limitations of this scoring method. It is worth noting that *HC Cons* does not perform any better than just counting up hotspots, but the combination of conservation and coevolution at hotspots (*KFC-E)* is a clear improvement over coevolution alone. This is likely because conservation does not have a strong signal on its own but serves to enhance the coevolutionary signal.

### Performance Comparisons for Four Evolutionary Scoring Functions

We inspected the efficiency in ranking the docked poses for four scoring functions: *HC Co-evol*, *HH Co-evol*, *NH-NH Co-evol* and *HC Cons,* for three complexes taken from the bound dataset (Fig 1). Although *HC Co-evol* and *HH Co-evol* use hotspot coevolution information to score poses, the *HC Cons* scheme uses the extent of conservation of hotspot residues and their contacts across the binding interface. The *NH-NH Co-evol* scoring function, however, employs the extent of coevolution of non-hotspot residues with their contacts in the binding partner. We verified the scoring trends using these four scoring functions for three complexes (PDB IDs: 1EP3, 3G5O, 1QOP) and found that *HC Co-evol* and *HH Co-evol* show similar performance in ranking the near-native poses higher than the non-native ones (Table S1). However, the performance was markedly reduced when non-hotspots and their extent of coevolution were used to score the docked poses.

*NH-NH Co-evol* did not identify a natively bound pose (i-RMSD ≤ 2 Å) in the top 10 scored poses for any of the three complexes analyzed (Table S1). Moreover, the funnel-like shape in the plot of score vs i-RMSD (Fig. 1) representative of the free-energy minimum and one the key characteristics of docking scoring functions, was absent for *NH-NH Co-evol.* However, the other three scoring functions using the KFC2-predicted hotspots consistently scored the near-native poses higher than the non-native ones and exhibited energy funnel-like behavior, emphasizing the greater importance of hotspots over non-hotspots in scoring the docked conformers accurately.

The performance of the *HC Cons* scoring scheme is comparable to that of *HC Co-evol* and *HH Co-evol* in that it produces the energy funnel and also detects near-native binding modes in the top scored poses (Fig. 1). However, *HC Cons* produces more false positives than *HC Co-evol* and *HH Co-evol*. Some of the non-native poses that have a lower score when using coevolution are ranked higher when conservation is used instead of coevolution (Table S1). A higher variance was observed for the i-RMSDs of the top 10 scored poses taken from *NH-NH Co-evol* and *HC Cons* compared to those taken from *HC Co-evol* and *HH Co-evol* (Fig. S1). We observed that for all the 53 complexes in the bound dataset, on average, *HC Co-evol* performed marginally better than *HH Co-evol*, correctly predicting three out of the top 10 poses scored by *HC Co-evol* as near-native structures. The performance was significantly worse for *NH-NH Co-evol*, identifying less than one near-native structure in the top 10 (Fig. S2). Thus, coevolution at predicted non-hotspots is not as effective as coevolution at predicted hotspots at identifying native binding models.

### Coevolution- and Conservation-based Scoring Function: *KFC-E*

Because some systems did better with a conservation-based score and some with a coevolutionary one, we propose a combined scoring function that utilizes both. *KFC-E* combines coevolution and conservation to score binding modes by considering the extent of conservation of hotspot residues and the coevolution of these residues with contacts at the interface of the binding partner. Specifically, it accounts for the strength of the coevolutionary signal across the interface (i.e., between hotspots and their contacts) relative to the signal within the interface of a binding partner (i.e., between the contacts themselves) (Fig. 2). If *i* is a hotspot residue on chain A and *j*_1_, *j*_2_ and *j*_3_ are residues in contact with *i* (i.e., within 7 Å of *i*), then *KFC-E* measures the cumulative coevolution of *i* with each of *j*_1_, *j*_2_ and *j*_3_ relative to the total coevolution between (*j*_1_, *j*_2_), (*j*_2_, *j*_3_) and (*j*_1_, *j*_3_). Our underlying hypothesis is that to maintain the binding interactions across the interface, the residues that are in contact with a given hotspot residue should co-evolve more strongly with the hotspot residue than among themselves. That is, the coevolutionary signal across the interface of the two binding partners should be stronger than the signal between residues forming the interface in either binding partner.

We compared the scoring performance between the *KFC-E* and *HC Co-evol* scoring functions for six complexes (PDB IDs: 1RM6 (chains A and B), 1TYG, 3G5O, 3MML, 4HEA (chains A and H), and 4HEA (chains J and K) (Fig. 3). The number of false positive hits in the top 10 poses was considerably lower for KFC-E than for *HC Co-evol* (Table S2). Moreover, the variance of i-RMSD for the top 10 scored poses for these six complexes is significantly higher for *HC Co-evol* compared to *KFC-E* (Fig. 4). The median i-RMSD for all the top 10 scored poses from *KFC-E* is also considerably smaller (1.8 Å) compared to that of *HC Co-evol* (3.1 Å). *KFC-E* identified only eight poses with i-RMSD ≥ 5 Å, whereas the number was much higher for *HC Co-evol* (28 poses). Thus, including information on hotspot conservation and the coevolution between the contacts filters out a substantial amount of false positives. The superior performance of *KFC-E* to that of *HC Co-evol* and the three other scoring functions (*HH Co-evol*, *NH-NH Co-evol*, *HC Cons)* was consistent for all 53 complexes in the bound dataset (Fig. 5) for all the top N categories. *KFC-E* has an average of more than three hits (i.e., poses with i-RMSD ≤ 2 Å) in the top 10 scored poses, whereas *NH-NH Co-evol* had the worst performance across all the top N categories with respect among the remaining four scoring functions.

**Figure 3.**
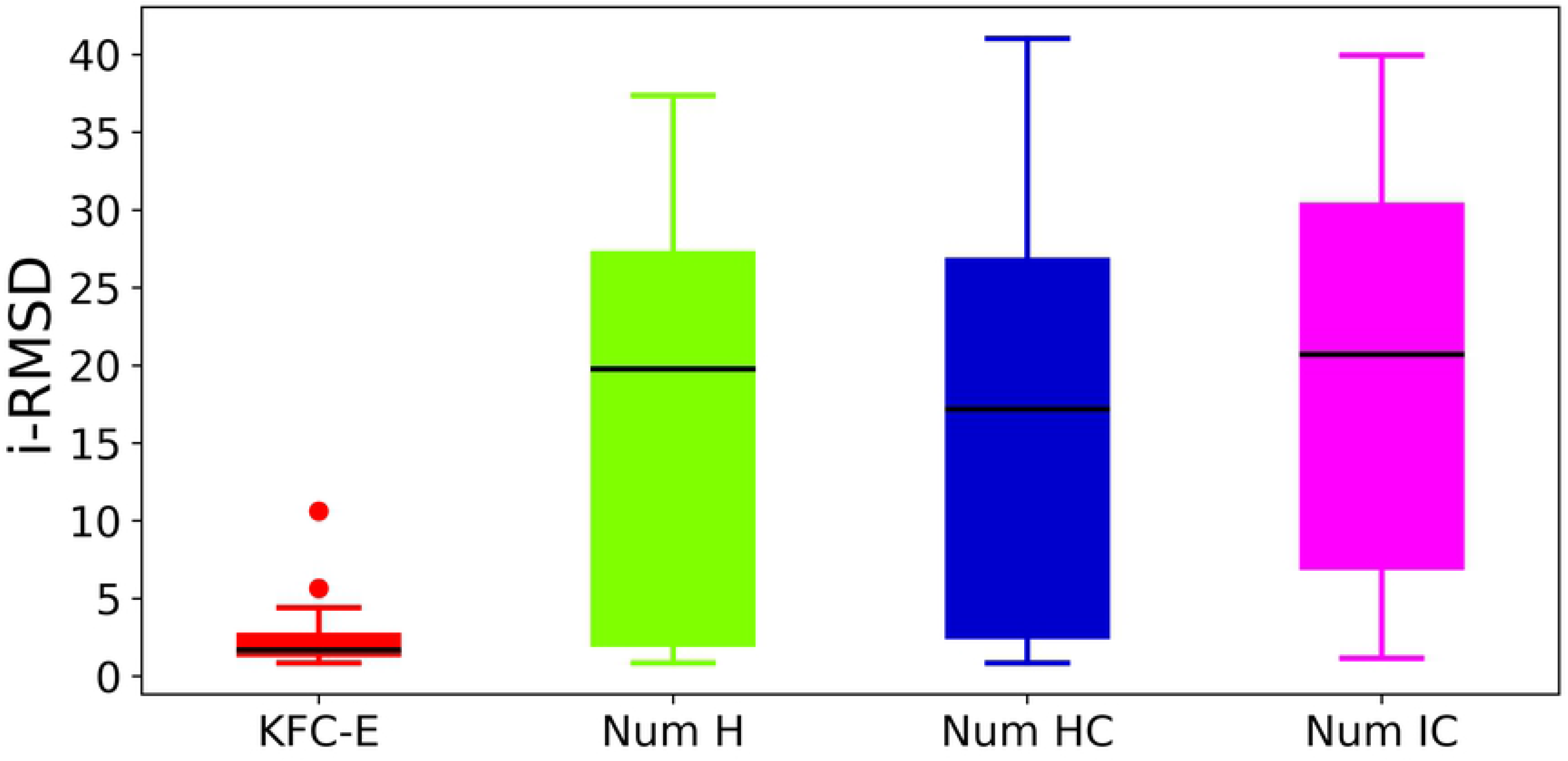
Comparison of trends between the *HC Co-evol* and *KFC-E* scoring functions on the bound dataset. Scoring trends for the *HC Co-evol* and *KFC-E* scoring functions for six complexes: 4-hydroxybenzoyl-CoA reductase (α and β subunits, PDB ID: 1RM6), thiazole synthase/ThiS complex (chains A and B, PDB ID: 1TYG), toxin-antitoxin complex (chains A and B, PDB ID: 3G5O), allophanate hydrolase complex (subunits 1 and 2, PDB ID: 3MML), respiratory complex 1 (chains A and H, PDB ID: 4HEA), and respiratory complex 1 (chains J and K, PDB ID: 4HEA). For each complex, the scoring trend for the *HC Co-evol* scoring function is shown on the left and that for *KFC-E* on the right.

**Figure 4.**
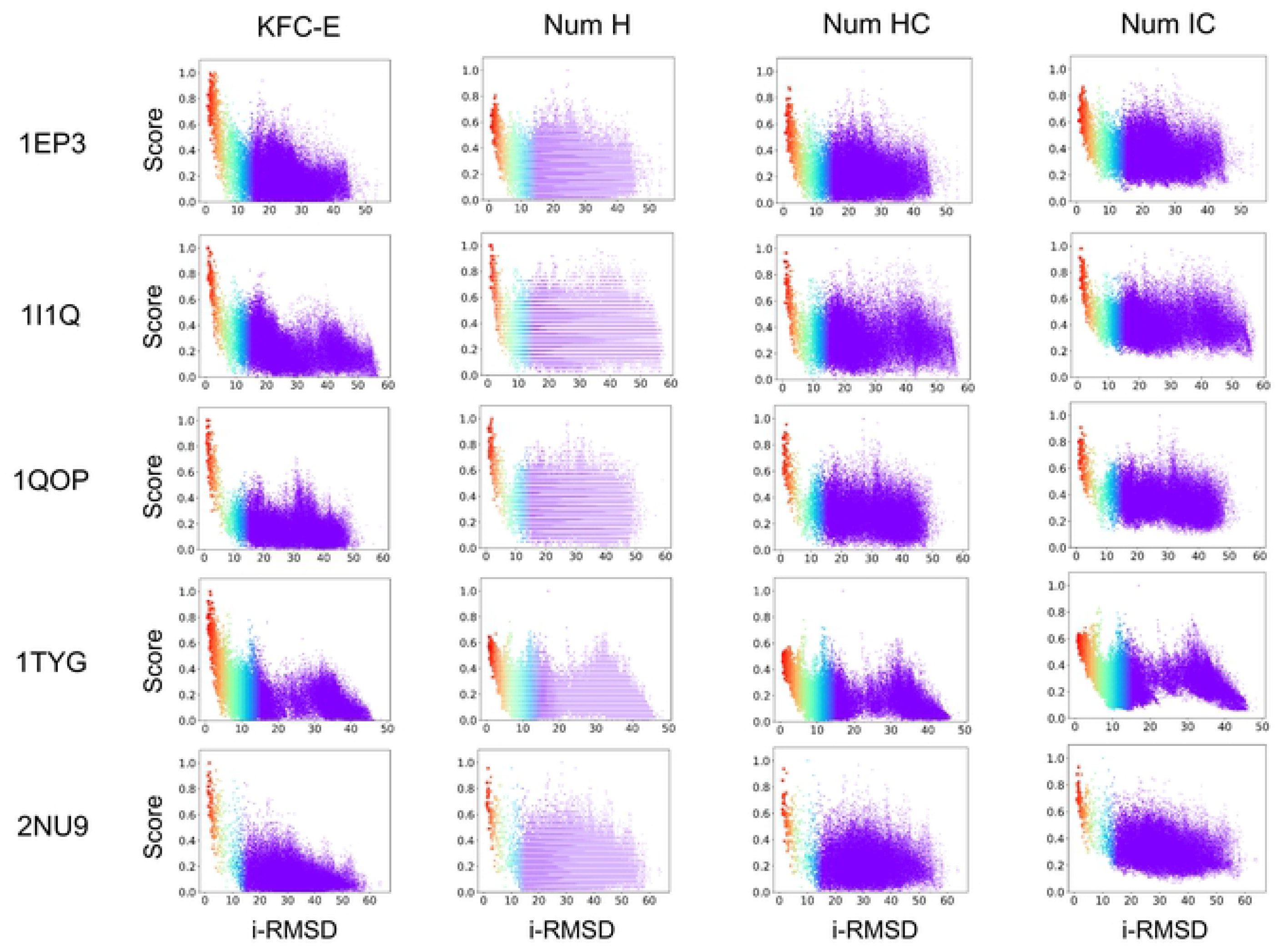
Distribution of i-RMSDs for the top 10 poses of six complexes scored with *HC Co-evol* and *KFC-E*. The calculations were performed on the same set of complexes as in Figure 3.

**Figure 5.**
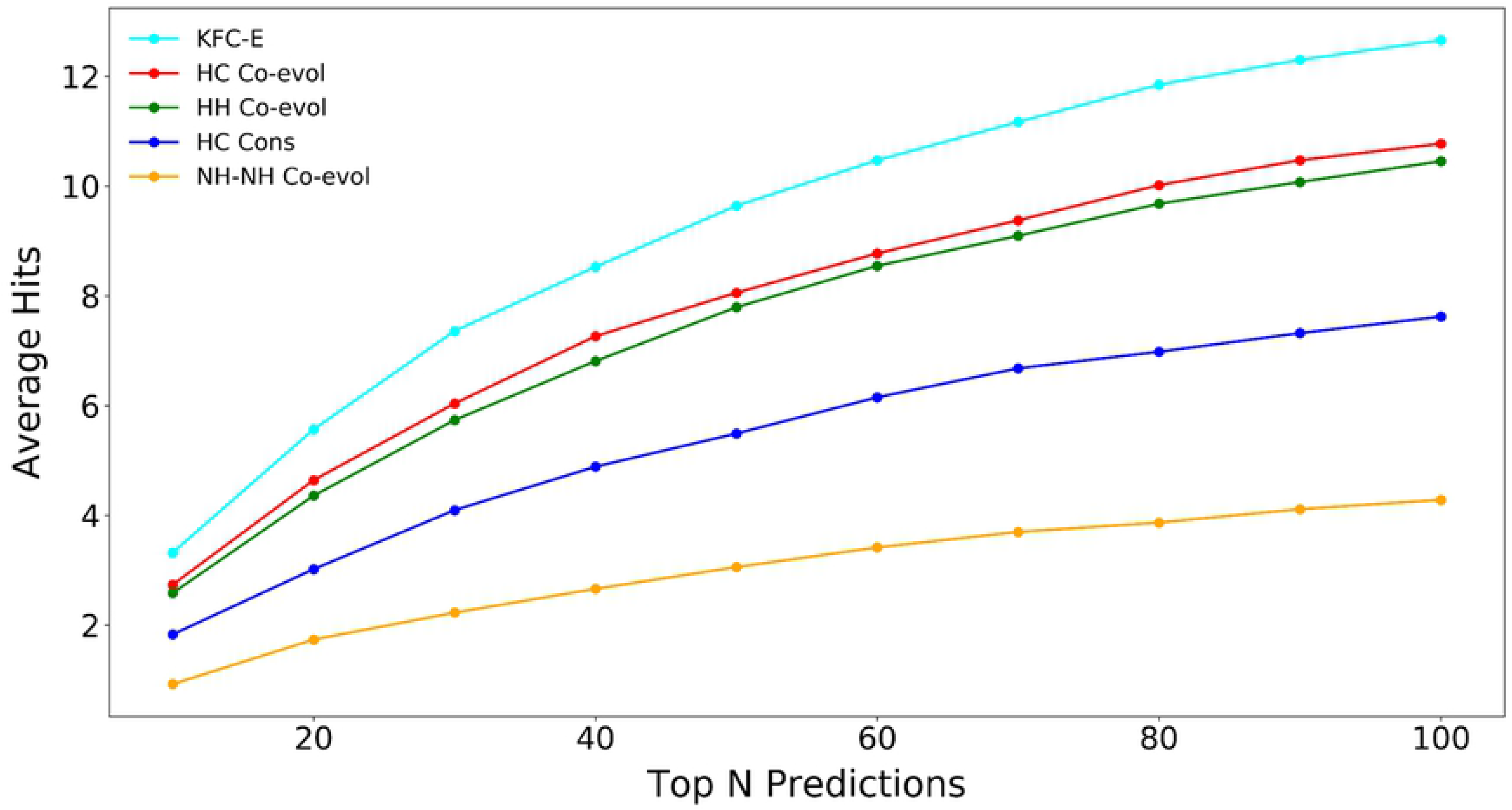
Average performance of *KFC-E* on the bound dataset. Average number of near-native hits in each of the top N categories for the *KFC-E* scoring function compared to the *HC Co-evol, HH Co-evol, HC Cons* and *NH-NH Co-evol* scoring functions.

### Performance of *KFC-E* versus Scoring Functions without Evolutionary Information

Does the inclusion of hotspot evolutionary information into a scoring framework make a difference in screening natively docked poses from non-native ones? How accurate are the results if we consider only the size of the binding interface, the number of hotspots, or just the number of hotspot contacts to score the docked poses? How do such scoring functions perform in scoring the docked poses compared to those that include evolutionary information? To answer these questions, we introduced three separate scoring functions whose scores correlate strongly with the size of the interface but do not include any evolutionary information. *Num H* calculates the total number of hotspot residues at the binding interface, while *Num HC* includes all contacts (i.e., distance between heavy atoms ≤ 7 Å) made by a hotspot residue with residues in the binding partner. *Num HC* is similar to *HC Co-evol* except that it does not include any coevolutionary information between the residues. Lastly, the scoring function *Num IC* estimates the size of the interface as the total number of unique contacts across the interface that are within 7 Å.

For each scoring function (*KFC-E*, *Num H*, *Num HC* and *Num IC*) we show the scoring trends for five complexes taken from the bound dataset (Fig. 6), with each row corresponding to the scoring results for an individual complex using each of the four scoring functions and each column describing the performance of a given scoring function for each of the five complexes. For each complex, *KFC-E* correctly identified a near-native pose and assigned it a high score. More importantly, it assigned low scores to non-native poses, thus reducing the number of false positives relative to the other three scoring functions. The median i-RMSD of the top 10 poses sampled by *KFC-E* is 1.7 Å (Fig. 7). In addition, the individual i-RMSDs (Table S3) for the *KFC-E* poses have considerably lower variance than those for the *Num H*, *Num HC* and *Num IC* scoring functions. The scoring functions that do not include any co-evolutionary information have a strong tendency to assign high scores to non-native poses and thus produce a high number of false positives. *KFC-E* identified only two poses with i-RMSD ≥ 5 Å in the top 10 scored poses of all five complexes (i.e., a total of 50 poses), whereas *Num H*, *Num HC* and *Num IC* identified 34, 36 and 44 such poses, respectively. The better performance of *KFC-E* is notable for the thiazole synthase/ThiS complex (PDB ID: 1TYG) for which none of the other three scoring methods identified a high-scoring natively bound complex. Thus, *KFC-E*, by virtue of its inclusion of evolutionary information, more accurately identifies native complexes and filters out false positives, a key aspect of scoring functions for protein-protein docking.

**Figure 6.**
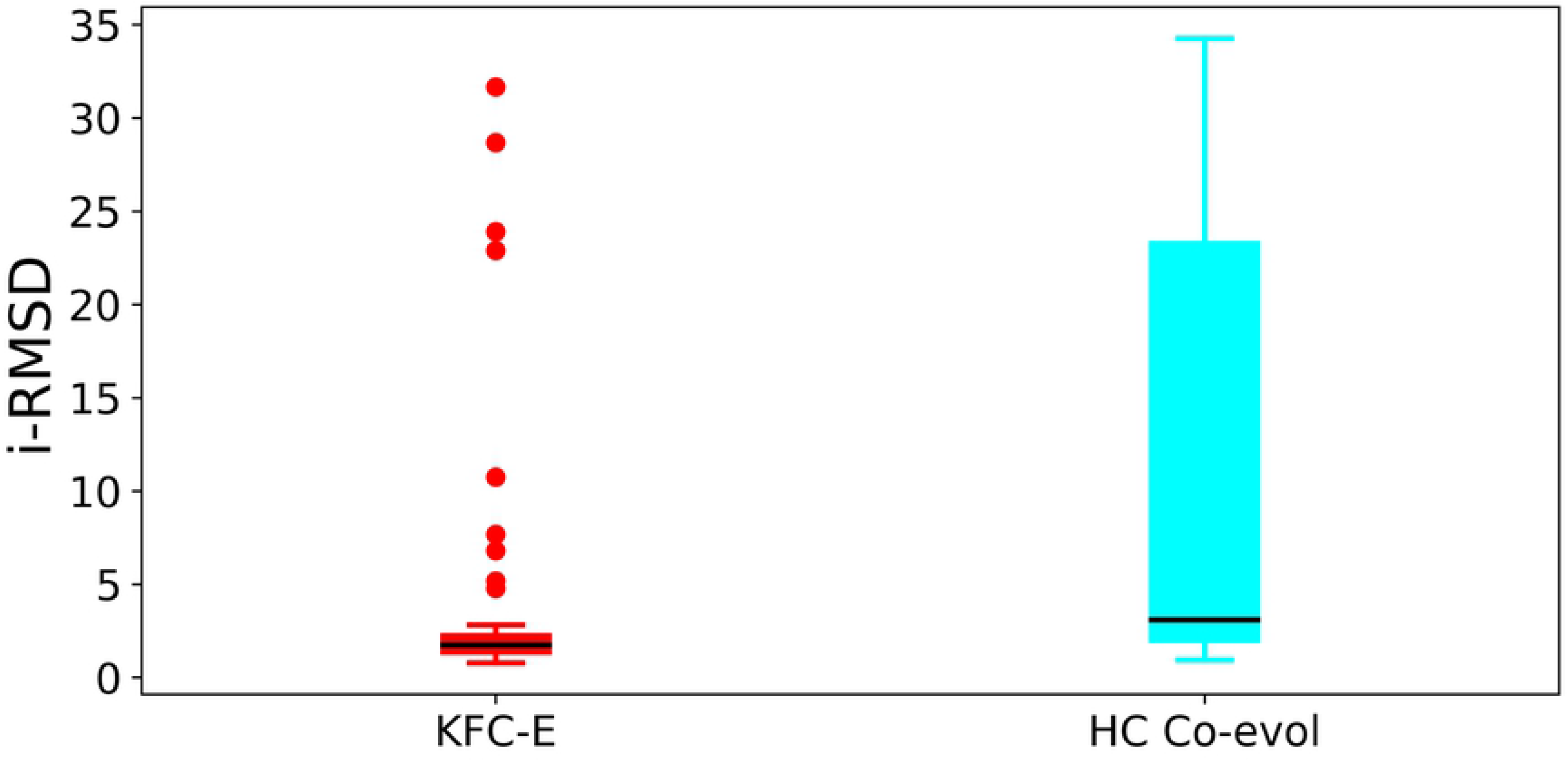
Comparison of *KFC-E* to the *Num H* (number of hotspots), *Num HC* (number of hotspot contacts) and *Num IC* (number of interface contacts) scoring functions. Each row shows the scoring trends for the four functions for five complexes taken from the bound dataset: dihydroorotate dehydrogenase (PyrD and PykR subunits, PDB ID: 1EP3), anthranilate synthase (components 1 and 2, PDB ID: 1I1Q), tryptophan synthase (α and β chains, PDB ID: 1QOP), thiazole synthase/ThiS complex (chains A and B, PDB ID: 1TYG), succinyl-CoA synthase (α and β chains, PDB ID: 2NU9).

**Figure 7.**
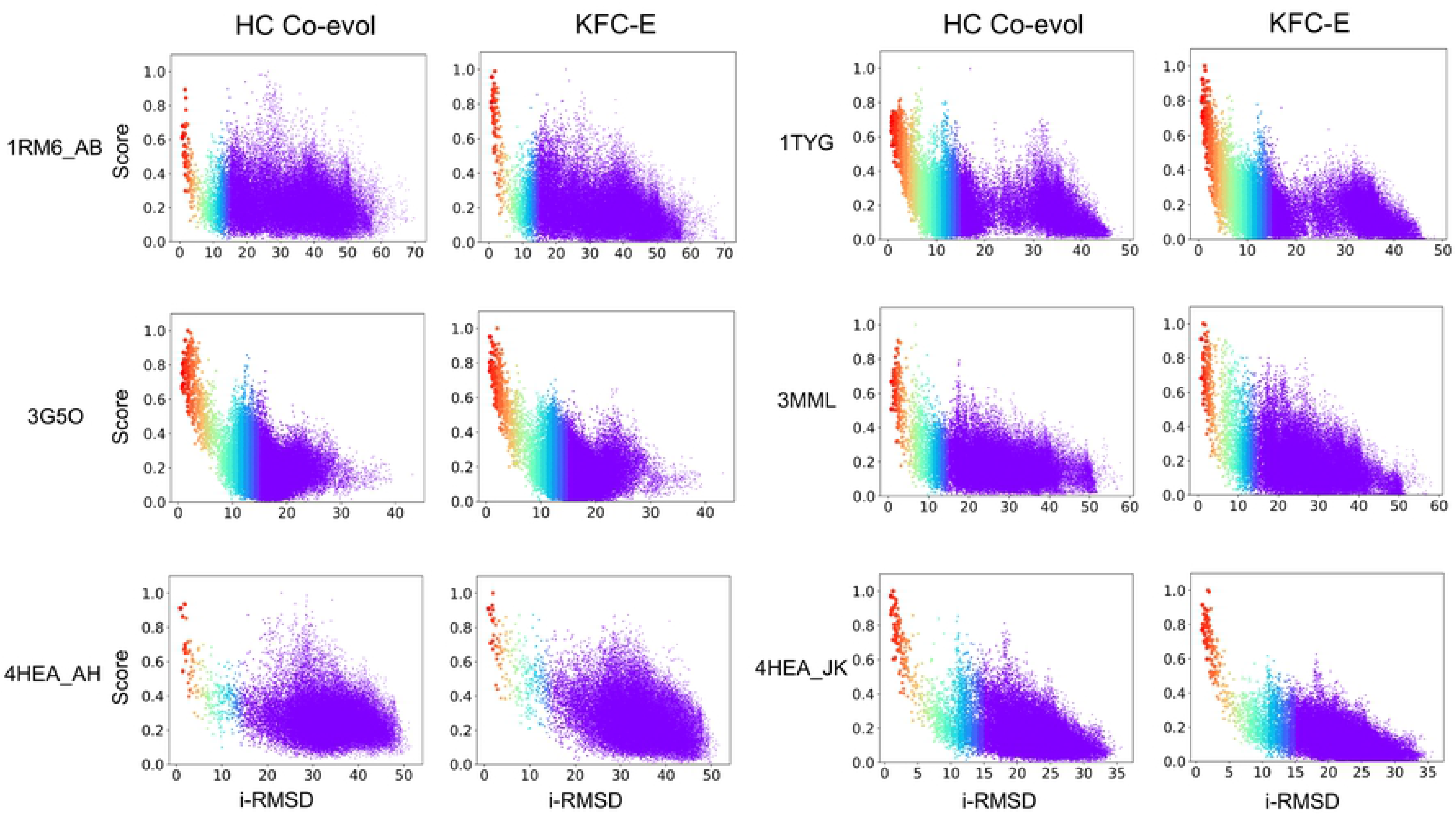
Distribution of i-RMSDs for the top 10 poses for 5 complexes scored with *KFC-E*, *Num H*, *Num HC* and *Num IC*.

We also examined the average performance of all eight scoring functions (see Table 3) for all 53 complexes in the bound dataset. *KFC-E* has a much higher number of average hits (near-native poses) than all other scoring methods tested (Fig 8A). The median i-RMSD of the top 10 scored poses of *KFC-E* is also lower (∼5 Å) compared to the remaining 7 scoring functions (Fig. S3). The performance of KFC-E on individual complexes is provided in Figures S4 and S5. Of the eight scoring methods, the three that use the coevolution information of KFC2a-predicted hotspots (*KFC-E*, *HC Co-evol* and *HH Co-evol*) show better performance than all other scoring methods.

**Figure 8.**
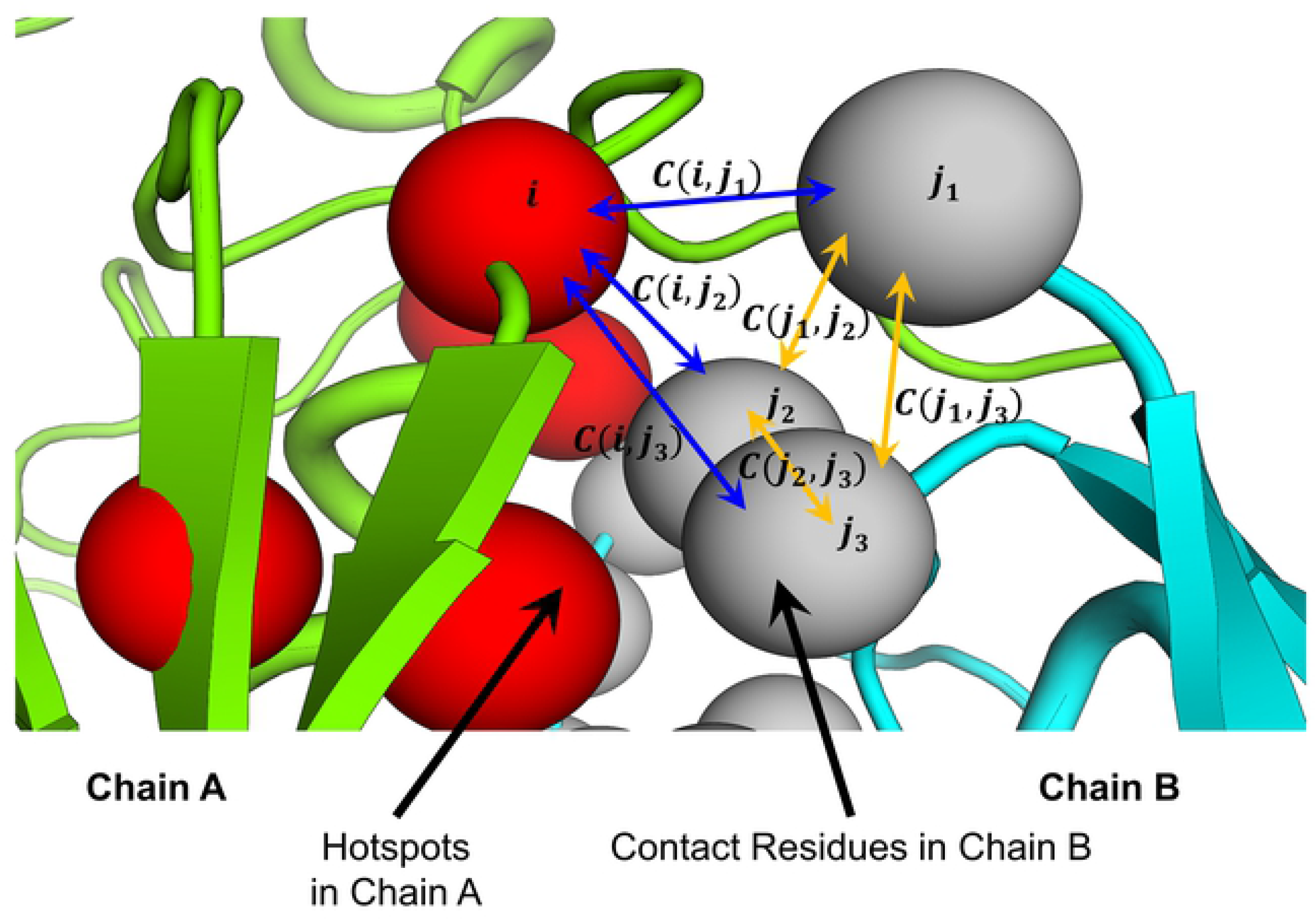
Performance of each scoring function assessed by the number and fraction of hits identified in each of the top N categories. (**A)** Average number of hits over all 53 complexes in the bound dataset in each of the top N categories. (B) Average fraction of hits identified by each scoring function in the bound dataset. For a given complex, the number of hits in the top N poses normalized by the total number of hits sampled by ZDOCK is shown. (**C**) The average fraction of hits for 46 complexes obtained after filtering out complexes for which ZDOCK sampled zero near-native binding modes. (**D**) Average fraction of hits for 25 complexes obtained after excluding respiratory complex 1 (PDB: 4HEA).

The only other scoring function included in our study that incorporates coevolutionary information is the *NH-NH Co-evol* function, which performs the worst and provides a low average number of hits. The superior performance of *KFC-E*, especially compared to *NH-NH Co-evol*, strongly suggests that not all the interface residues drive the evolution of bound complexes. Rather, it is specifically the hotspot residues that are most important, and the coevolutionary signal captured using hotspot residues is stronger than that from non-hotspot residues. This performance gain also attests to the accuracy of KFC2a in making reliable predictions for hotspot residues.

We also considered the average fraction of hits identified in each of the top N prediction categories (Fig. 8B) and consistently observed superior performance with *KFC-E* scoring method. Of the 53 complexes, only 46 had at least one near-native pose sampled by ZDOCK. We repeated our calculations for the average fraction of hits in each of the top N categories for the 46 complexes (Fig. 8C) and observed the same trend as before, but with an increase in the average fraction of hits. For these 46 complexes, the average number of near-native hits in the top 10 *KFC-E* poses was four, which is almost twice as many as the scoring methods using only conservation or no evolutionary information of hotspot residues (Fig. S6). We verified the number of near-native hits identified in the top 1 and top 5 ranked poses by *KFC-E* (Table 4) and found that for 38% of the complexes (20/53), *KFC-E* identified a near-native pose as the top-ranked and for 55% of the complexes (29/53), it identified near-native poses in the top 5. The performance of *KFC-E* was worse for the respiratory complex (4HEA) compared to all the other complexes. Of the 21 4HEA complexes that had at least one sampled near-native pose, *KFC-E* identified a near-native binding mode in the top 10 for only eight cases (Fig. S5). Upon excluding 4HEA, *KFC-E* identified an average of ∼22% of the total sampled near-native poses in the top 10 scored poses (Fig. 8D) and ∼6 of the top 10 poses identified by *KFC-E* are near-native (Fig. S7).

### Performance of *KFC-E* on an Unbound Dataset

Bound protein complexes resolved with methods such as X-ray crystallography usually have good shape complementary between monomers, which is often achieved through conformational changes of the binding interfaces upon binding. Consequently, pulling apart and re-assembling the interacting protein subunits is generally considered an easier problem in protein-protein docking because the subunits are already present in their optimal binding conformations. A more challenging problem is assembling the subunits of two proteins that are known to interact but with an unknown binding mode. In the present study, we considered 17 unbound complexes (Table 2) compiled from the benchmark dataset (see Methods). We used previously identified orthologous protein structures as templates to generate homology models for one or both binding partners in each complex. These models were then used as substitutes for the binding partners, mimicking a situation in which the subunits to be docked are structurally different from those in the natively bound form and therefore presenting a more challenging problem for sampling the binding modes as well as scoring them. We followed the same procedure as for the bound dataset, i.e., sampling binding modes with ZDOCK and then scoring the modes with *KFC-E*. Because the use of homology models can introduce structural differences with respect to the native complex, we opted for a more permissive i-RMSD cutoff of 3 Å to define near-native poses.

Using the 3 Å i-RMSD cutoff, *KFC-E* identified near-native poses in the top 10 for 4/7 complexes (1EP3_AB, 1RM6_BC, 3PNL_AB, 3RRL_AB) that had at least one near-native pose sampled by ZDOCK (Table 5). Among the three cases for which it failed to identify any native bound poses in the top 10 (1QO_AB, 1W85_AB, 3IP4_AB), for 1QOP_AB the pose with the lowest i-RMSD ranked in the top 10 was conformationally similar to the native bound form but had i-RMSD = 4.1 Å. In Figure 9E, the smaller subunit (colored in cyan) differs from the native pose primarily in the conformation of a loop at the interface, while the remainder of the docked structure closely resembles that of the native form. For two complexes (1B70_AB and 1I1Q_AB), *KFC-E* identified the lowest i-RMSD structure sampled by ZDOCK in the top 10 ranked poses. Figure 9 shows the best pose (lowest i-RMSD) in the top 10 poses identified by *KFC-E* for seven complexes in the unbound set. Plots of the trends in *KFC-E* scores for each complex in the unbound set are provided in Fig. S8.

**Figure 9.**
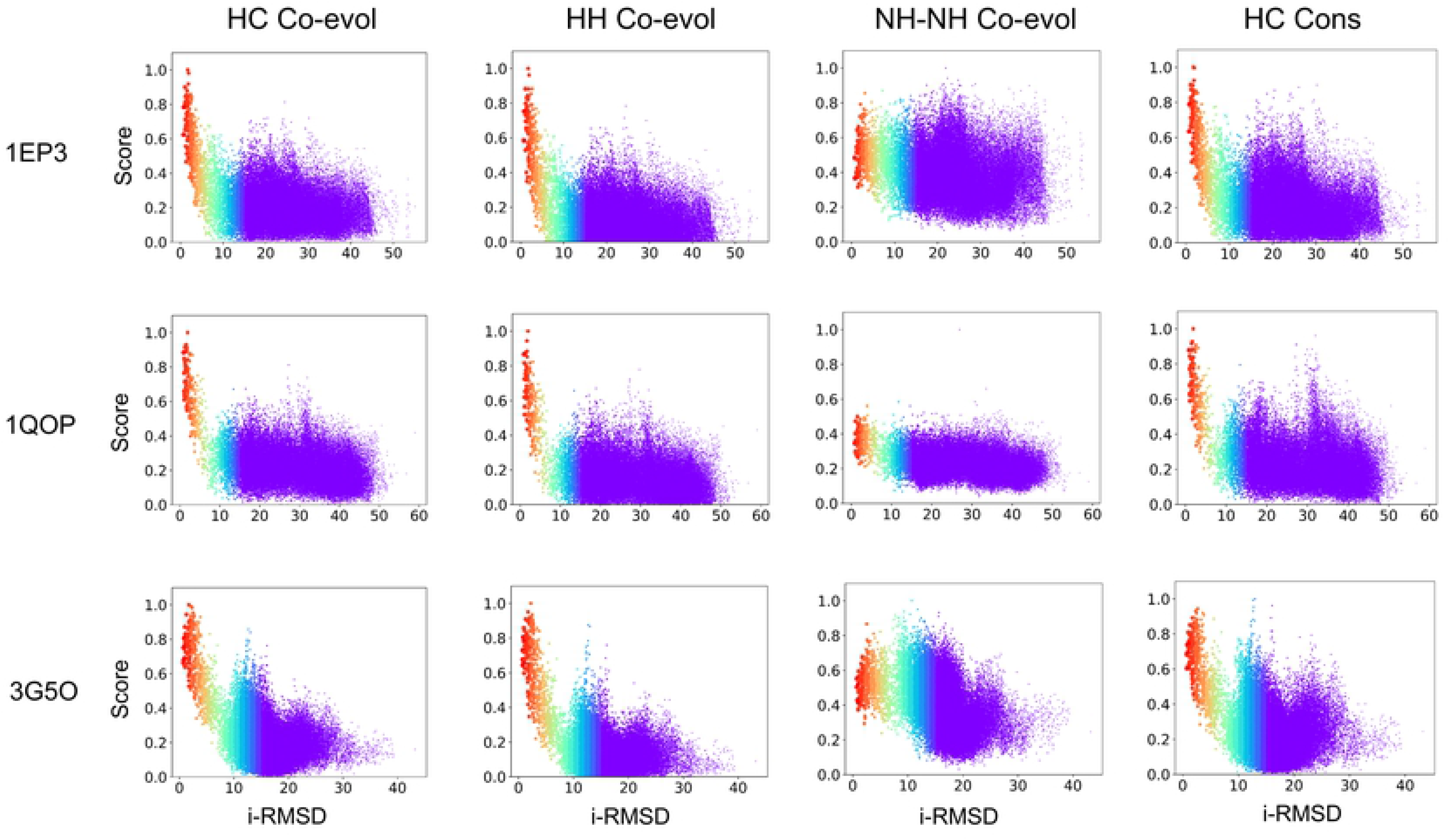
Lowest i-RMSD pose from the top 10 *KFC-E* poses in the unbound dataset. The complexes included are: (**A**) phenylalanyl t-RNA synthetase (chains A and B, PDB ID: 1B70). (**B**) anthranilate synthase (chains A and B, PDB ID: 1I1Q), (**C**) 3-oxoadipate CoA-transferase (chains A and B, PDB ID: 3RRL), (**D**) 4-hydroxybenzoyl-CoA reductase (chains B and C, PDB ID: 1RM6), (**E**) tryptophan synthase (α and β chains, PDB ID: 1QOP), (**F**) DHA kinase (chains A and B, PDB ID: 3PNL), and (**G**) dihydroorotate dehydrogenase (PyrD and PykR subunits, PDB ID: 1EP3). In each case, the native complex is shown in gray and the subunits from the pose are shown in green and cyan.

We also compared the performance of KFC-E to the raw ZDOCK score in identifying near-native hits (Table S4 and Fig. S9) and found that for 2/7 cases above that ZDOCK identified a near-native structure in its top 10 (Table S4). This was true twice as often for *KFC-E*, which returned a lowest iRMSD or near-native structure for 4 of 7 examples. Two additional examples returned structures with iRMSD 4.1 Å and 5.1 Å in the top 10 *KFC-E* scores.

### Limitations of the *KFC-E* Scoring Function

Because *KFC-E* uses residue-residue coevolution to score poses, its predictions depend on the quality and accuracy of the paired alignments used for generating the coevolutionary contact matrices as well as on the accuracy of the predicted residue-residue coevolution scores. Another key aspect is the number of aligned sequences required for the accurate prediction of co-evolving contacts. Though previous studies have indicated the necessity of having at least as many sequences as the combined length of the interacting proteins to predict protein-protein contacts accurately [39], others have shown that the coevolutionary signal captured with fewer sequences (i.e. < 100) can predict protein-protein complexes [11,40]. We used previously generated paired alignments with large number of sequences, which ensures a high signal-to-noise ratio for coevolution. Because we also coupled the coevolution with information on hotspots and their conservation, it is possible that accurate docked poses could be obtained with alignments containing fewer sequences, but that remains to be tested. The limited number of examples for which we have reliable data on orthologs points to the need for better methods for protein functional annotation.

## Conclusions

Using hotspots predicted by KFC2a, we have developed a method that includes evolutionary information of hotspots to score docked poses of protein-protein complexes. Our observation that the combination of coevolution and conservation can successfully predict near-native protein complexes is supported by past work on contact maps prediction [41,42]. Because hotspots contribute significantly (> 2 kcal/mol) to the binding free energy of protein-protein complexes, they should either be under higher evolutionary pressure than other interface residues. Our scoring function, *KFC-E*, capitalizes on the availability of large number of protein sequences to capture residue-residue coevolution and residue conservation, and estimates the strength of evolutionary signals across the interface of protein complexes. Our study suggests that for the present dataset, hotspot coevolution is a better predictor of native protein complexes than hotspot conservation. However, combining the two into a single scoring function (*KFC-E*) gives the best performance overall. We have tested our method on a benchmark set of bacterial protein complexes, but its performance remains to be tested on eukaryotic proteins for which correctly pairing the orthologs is a major challenge. Recent contributions however, shed light on how to generate paired alignments for eukaryotes for such coevolutionary studies and may be of significance [43–45] for performing the coevolution calculations.

## Acknowledgements

SJC was supported by NIH/NIGMS-IMSD Grant No. R25GM086761 and a National Science Foundation Graduate Research Fellowship under Grant No. (2017219379). JMP, JCM and SKM were supported by the Laboratory Directed Research and Development program at Oak Ridge National Laboratory, which is managed by UT-Battelle, LLC, for the DOE under Contract No. DE-AC05-00OR22725. This work used resources of the Compute and Data Environment for Science (CADES) at ORNL, which is supported by the Office of Science of the U.S. Department of Energy under Contract No. DE-AC05-00OR22725. The authors would like to thank Sergey Ovchinnikov for sharing the dataset of bound and unbound protein-protein complexes.

## Author Contributions

SKM and JCM designed the research. SKM and SJC conducted research. SKM, SJC, JMP and JCM equally contributed to writing the paper and analyzing the results. JCM supervised the research.

